# Whole Metagenome Sequencing: not Deep Enough for Complete Microbial Function Recovery

**DOI:** 10.1101/2025.11.04.685665

**Authors:** Jia Liu, Modupe Oluseun Coker, Nosayaba Osazuwa-Peters, Oluwaseun Peter, Nosakhare Lawrence Idemudia, Nicolas F. Schlecht, Ozoemene Obuekwe, Fidelis E Eki-Udoko, Yana Bromberg

**Affiliations:** Department of Biology, Emory University, Atlanta, Georgia, USA; Department of Basic and Translational Sciences, School of Dental Medicine, University of Pennsylvania, Philadelphia, PA; Center for Clinical and Translational Research, School of Dental Medicine, University of Pennsylvania, Philadelphia, PA; Department of Epidemiology, Geisel School of Medicine at Dartmouth, Hanover, New Hampshire; Department of Biostatistics, Epidemiology and Informatics, School of Medicine, University of Pennsylvania, Philadelphia, PA; Department of Head and Neck Surgery & Communication Sciences, Duke University School of Medicine, Durham, North Carolina; Department of Population Health Sciences, Duke University School of Medicine, Durham, North Carolina; Duke Cancer Institute, Duke University, Durham, North Carolina; Duke Global Health Institute, Durham, North Carolina; Institute of Human Virology Nigeria, Abuja, Nigeria; Medical Laboratory Services, University of Benin Teaching Hospital, Benin City, Nigeria; Department of Cancer Prevention and Control, Roswell Park Comprehensive Cancer Center, Buffalo, New York; Department of Oral and Maxillofacial Surgery, School of Dentistry, University of Benin Teaching Hospital, Benin City, Nigeria; Department of Child Health, University of Benin Teaching Hospital, Benin City, Nigeria; Department of Computer Science, Emory University, Atlanta, Georgia, USA

**Author notes:** Corresponding authors: Jia Liu, Yana Bromberg.

**Keywords:** whole metagenome shotgun sequencing, function, read depth, batch effects

## Abstract

**Background:** Whole metagenome shotgun sequencing (WMS) is widely used to profile microbial function. However, technical variability in sequencing and analysis often obscures true biological patterns. Large-scale studies are particularly susceptible to batch effects, such as differences in sequencing depth and platform and annotation strategies, as well as sample-to-flow-cell assignments. However, the relative effects of these factors on functional inference in such studies have yet to be systematically evaluated.

We analyzed oral-rinse WMS data from a study cohort including 671 Nigerian youths aged 9-18, sequenced on two Illumina platforms. Microbial molecular functionality encoded in these data were annotated using the mi-faser/Fusion pipeline, to capture the broad functional repertoire, and HUMAnN 3/EC numbers pipeline to characterize curated enzymatic activities. We then quantified how technical factors and batch effects shaped the recovery of microbial functionality.

**Results:** Three findings of our work were most salient. First, we observed that the choice of annotation strategy traded off between breadth and specificity of functional coverage. Second, we found that low-prevalence functions were disproportionately lost at shallow sequencing depths, indicating that in e.g. case-control studies with few representatives of the minor class, sequencing depth could critically impact study resolution. Finally, using our newly developed model relating sequencing depth to functional recovery, we demonstrated that increasing sequencing depth does not directly or proportionally improve functional recall. That is, at as little as 10% of this study’s sequencing depth, 30% of the estimated complete microbiome functional repertoire was detectable. However, even at the full depth used in this study, we were only able to recover an estimated 60% of that complete functional repertoire.

**Conclusions:** Together, these findings and our depth-to-function mapping framework provide practical guidelines for the design and interpretation of WMS studies. Coordinating sequencing depth planning with annotation strategy, experimental design, and rigorous batch control is thus essential for robust detection of microbial functions and for ensuring reproducible microbiome insights.

## Background

Whole metagenome shotgun sequencing (WMS) enables comprehensive profiling of microbial communities, capturing both taxonomic composition and functional potential directly from sample DNA. Functional profiling, in particular, allows researchers to identify microbial genes and pathways specific to host groups or environments, providing insight into microbial contributions to health, disease, and ecological adaptation^1-4^. While functional differences across samples are often interpreted as biologically meaningful, many technical and experimental factors, e.g. sequencing depth, batch effects, and functional annotation strategies, also contribute to variation, thereby confounding identification of phenotype drivers.

Sequencing depth is a critical determinant of functional recovery, governed primarily by two factors: (1) functional diversity of the sample’s microbial community and (2) fraction of microbial DNA within the sample. An environmental or host-associated sample represents only a subset of the true community, and WMS further samples only a fraction of the DNA molecules present in that sample (**Figure 1**). Sequencing depth determines how many of these DNA molecules are captured, and thus how much of the underlying functional diversity is potentially recovered.

**Figure 1:**
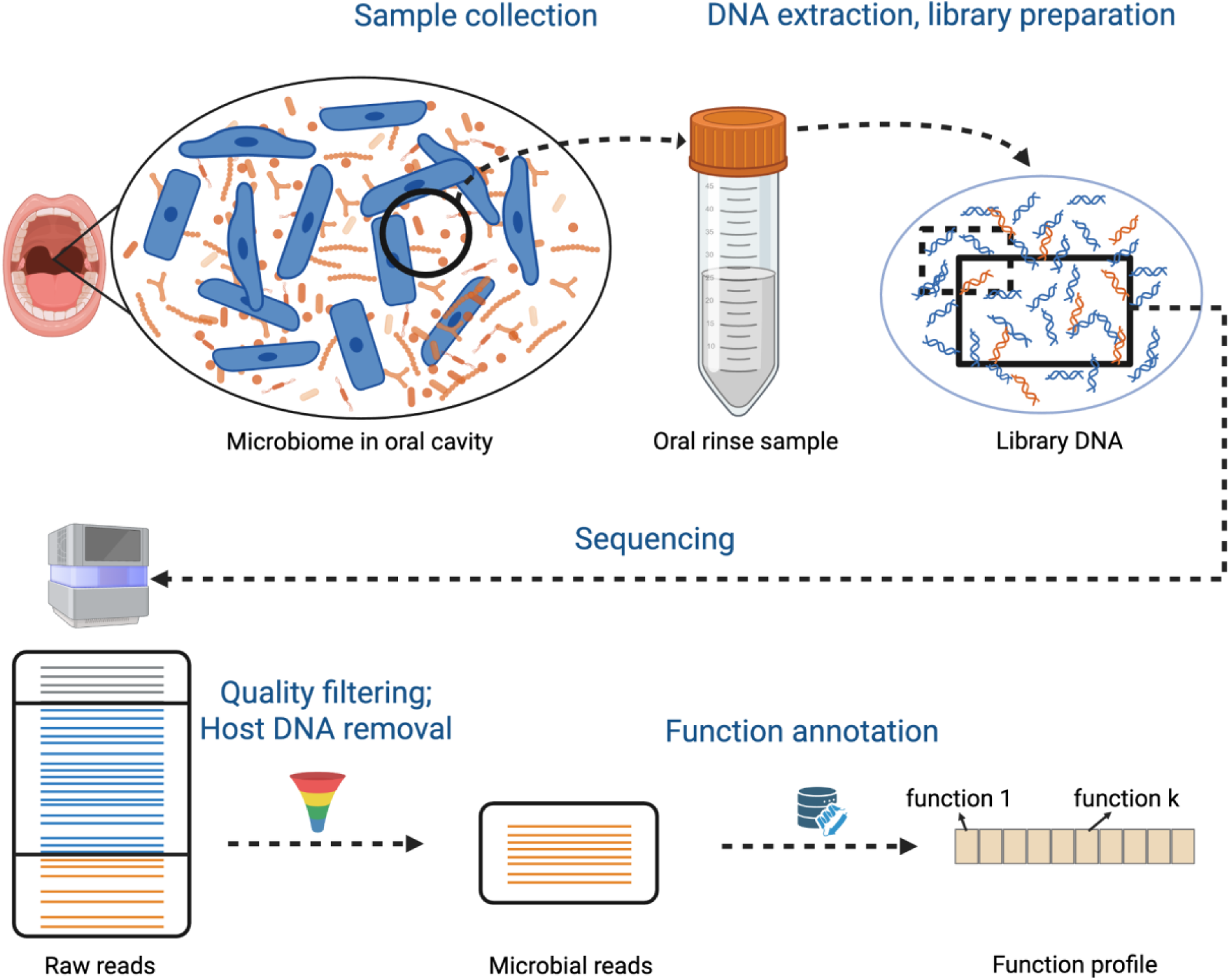
Oral microbiome sampling, sequencing, quality assurance, and functional annotation. The oral microbiome of each of the youths in this study was sampled using the oral rinse procedure (bold black circle in oral cavity microbiome). DNA was extracted from these samples, capturing host (blue) and microbial (orange) DNA in varying proportions. The fraction of DNA that was sequenced (black square) resulted in DNA reads, which we labeled as low-quality (grey) or identifiable host reads; the rest were deemed microbial. Microbial reads were further annotated via Fusion or EC numbers to generate individual-specific functional profiles.

Given that the true functional diversity in a natural (non-simulated) microbiome sample is typically unknown, the sequencing depth necessary or sufficient for describing this functionality is also uncertain. Prior work has shown that depth strongly impacts detection of low-abundance features such as rare taxa or functions^5-7^. This challenge of estimating the “right” sequencing depth for capturing the underlying microbial functional diversity is further complicated by host DNA contamination, which can dominate host-associated environments like the oral cavity^8^.

Beyond sequencing depth, other technical factors also confound biological signals, particularly in large scale studies or meta-analyses that integrate datasets from diverse sources, such as the Integrative Human Microbiome Project (iHMP)^3^, MGnify^9^, and MetaSUB^10^. Samples may be processed in separate batches (either randomly or deliberately assigned), sequenced across multiple flow cells, or annotated via different strategies, etc. These sources of heterogeneity can affect the number and types of microbial functions detected, potentially confounding biological interpretations even in the absence of true biological differences^7, 11^. Previous studies have explored the impact of methodological factors such as sequencing depth, DNA extraction, library preparation, and downstream analytical pipelines on functional composition^5-7, 11-13^. While these studies have provided valuable insights, they have been limited by small sample size or use of simulated data; hence, our understanding of how technical factors influence microbiome annotation remains limited.

Here, we set out to address two key questions: (1) to what extent does WMS capture the true functional diversity present in oral microbiome samples, and (2) how do technical decisions shape this recovery?

## Methods

### Data preparation

Oral rinse samples were collected from 671 participants (ages 9-18 years) at the baseline clinical visit of a longitudinal study, Human Papillomavirus, Human Immunodeficiency Virus, and Oral Microbiota Interplay in Nigerian Youth (HOMINY)^14^. Of these, 445 were either with perinatally acquired HIV or exposed in utero but uninfected. The remaining 226 participants were not exposed to, and therefore uninfected with HIV; designated healthy in this study. All 671 samples underwent whole metagenome shotgun sequencing (WMS)^14^.

Sequencing was carried out in two batches:

● Batch 1: 544 initial-visit samples were sequenced on an Illumina NovaSeq 6000 platform with 150 bp paired-end (PE150) reads. Samples were distributed across three S4 flow cells in the order of participant enrollment: (1) 182 samples on flow cell fc_2310, (2) 182 on fc_2311, and (3) 180 on fc_2401. Each S4 flow cell on NovaSeq 6000 has an expected throughput of approximately 8.0-10.0 billion PE150 read pairs, corresponding to an expected per-sample yield of 44.4-55.6 million read pairs given the number of samples per cell (180-182).
● Batch 2: The remaining 127 initial-visit samples were sequenced together with 606 follow-up visit samples (not analyzed in this study), for a total of 733 samples. Sequencing of PE150 reads was performed on a 25B flow cell of the Illumina NovaSeq X platform. The expected throughput for a 25B cell is approximately 26.7 billion paired end read pairs, which, when distributed across 733 samples, corresponds to an expected yield of 36.0 million read pairs per sample.

Raw sequencing reads were demultiplexed using bcl-convert^15^ (v3.8.2-12-g85770e0b). Quality filtering was performed using fastp^16^ (v0.23.1) to remove reads with low average quality scores (“-e 25”) and short lengths (“-l 70”). Note that we identified no publications that would define standard parameters to be used for this filtering; thus, our decisions were based on previous studies^17, 18^. The filtered reads were then aligned to the human reference genome (GRCh38.p14) with BWA-MEM^19^ (v0.7.17) using default parameters to remove host DNA contamination. Paired-end reads, for which neither mate mapped to the human genome, were retained as microbial and used for downstream functional annotation.

### Fusion/mi-faser function annotation

The Fusion database^20^ comprises 433,891 microbial functions, spanning both well-characterized and putative functions, curated from 8,906 distinct, fully assembled bacterial genomes. The quality-filtered, host-depleted microbial reads were annotated with Fusion functions using mi-faser^21, 22^ (v1.61). The counts of reads per Fusion function were reported as functional abundance profiles for each sample.

### EC Number/HUMAnN 3 function annotation

To generate enzymatic functional profiles, we used the HUMAnN 3^23^ (v3.9) pipeline. Quality-filtered and host-depleted microbial reads were first aligned against species’ pangenomes to get functional annotations. Functions of unmapped reads were established by alignment against the UniRef90 protein database^24^ (version 201901b). The resulting gene family abundance tables were regrouped to EC numbers^25^ using the HUMAnN 3 utility script humann_regroup_table.

### Generation of relative abundance profiles

For both Fusion- and EC number-profiles, the functions with read counts <5 were removed to reduce the influence of potential annotation errors. Counts were then normalized by the total number of microbial reads per sample to generate relative abundance profiles for downstream comparisons.

### Sub-sampling to simulate reduced microbial read depth

To evaluate the impact of microbial read depth on functional profiling, we simulated reduced-depth conditions by randomly sub-sampling without replacement 10%, 30%, 50%, 70%, and 90% of microbial read pairs from each of the 671 samples. To avoid contamination and confounding, sub-sampling was performed after quality filtering and host DNA removal, using only microbial read pairs. For all simulated samples, Fusion relative abundance functional profiles were generated as described above. For the 70% sub-sampled sets, EC number-based relative abundance profiles were also generated as described earlier.

### Fusion functional dropout analysis at 70% sub-sampling

We assessed how function abundance and prevalence influenced dropout at a lower simulated sequencing depth (70% sub-sampling, i.e. 30% microbial read reduction).

For *abundance* dropout evaluations, Fusion functions within each original sample were ranked by relative abundance and assigned to decile bins (0-10%, 10-20%, …, 90-100% of each sample). A function was defined as “lost” if it was absent from the corresponding sub-sampled profile. The percentage of lost functions was calculated for each abundance bin in each sample.

For *prevalence* dropout evaluations, we first computed each Fusion function’s prevalence across the 671 original samples, i.e. the number of samples in which it was detected at full sequencing depth. We then calculated the proportion of those samples where the function was lost after sub-sampling (dropout rate).

Pearson and Spearman correlations were used to quantify the relationship between prevalence and dropout rate.

### Saturation curve modelling

To quantify the relationship between microbial read depth R, expressed as the number of read pairs, and the number of detected Fusion functions F, we evaluated three models: Michaelis-Menten^26, 27^ (Eqn. 1), Weibull growth^28^ (Eqn. 2), and the Hill equation^29, 30^ (Eqn. 3),

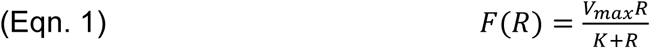

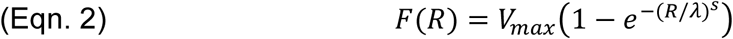

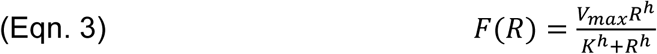

where *R* is microbial read pairs, V_max_ is the asymptotic plateau of recoverable functions, K is the half-saturation constant representing the microbial reads required to achieve half of V_max_, h is a slope/shape parameter (with h=1 reducing the Hill model to Michaelis-Menten), and λ (scale) and s (shape) are Weibull parameters controlling the depth scale and concavity, respectively.

For each model, we first fit a ‘pooled’ curve, using all samples jointly across all depths. We evaluated model performance using coefficient of determination (*R*²), root mean squared error (RMSE), and Akaike Information Criterion (AIC) measures. We then fit per-sample curves and evaluated their ability to capture our data using two holdout strategies: leave-top-depth-out (leave 100% complete dataset out) and leave-bottom-depth-out (10% subsampled dataset withheld). In both cases, we computed residuals (predicted minus observed function count) on the held-out point and summarized bias (median residual) and inter quartile range (IQR).

Across both pooled and per-sample fits, the Hill model achieved the best overall fit (**SOM_Table_1, SOM_Fig.1-2**) and was therefore selected for downstream analyses. After model selection, we re-fit the Hill model to all available sequencing depths for (1) each sample individually, and (2) over all samples jointly, to obtain the final per-sample and population-level parameter estimates (V_max_, K, h).

From these curves, we estimated the proportion of functions recovered at each sample’s full microbial read depths relative to the plateau. We also used the curves to interpolate and extrapolate the microbial read depths required to achieve specified function retention thresholds (50-95% of the plateau).

Corresponding raw read requirements for each microbial read estimate were derived by scaling microbial read count using the median microbial-to-raw read ratio across all 671 samples.

### Evaluation of Flow-cell effects

To assess the potential technical batch effects of sequencing flow cells, we analyzed Fusion-annotated relative functional abundance profiles from 671 oral rinse samples sequenced across four flow cells (NovaSeq 6000 S4: fc_2310 n=182, fc_2311 n=182, fc_2401 n=180; NovaSeq X 25B: fc_b2 n=127). To minimize clinical variation and better isolate technical effects, analyses were restricted to the 226 healthy participants (fc_2310: n=22, fc_2311: n=90, fc_2401: n=59, fc_b2: n=55; total: N=226, **SOM_Table_2**).

Functional profiles for these 226 healthy samples were represented as vectors of relative abundances of Fusion functions. Each vector length was equal to the size of the union of all functions annotated across all 226 samples (57,061 functions total). Each absent function per profile was set to zero.

To quantify differences in community composition, for all pairs of function profiles we calculated (1) Bray-Curtis dissimilarity, capturing abundance-weighted differences and (2) Jaccard dissimilarity, capturing presence/absence differences. Each dissimilarity matrix was then used as input to permutational multivariate analysis of variance (PERMANOVA; 9,999 permutations) to test whether microbial functional composition differed among flow cells. Here, a significant result would indicate that differences in functional profiles are attributable, at least in part, to whether the samples in a given pair come from different sequencing flow cells. If the PERMANOVA result was significant, we further conducted pairwise PERMANOVA on Bray-Curtis and Jaccard dissimilarity matrices among all flow cell combinations, with p-values adjusted for multiple testing using the Benjamini-Hochberg procedure.

## Results

We analyzed oral rinse microbiome WMS data (**Figure 1**) from youth participants collected at the initial visit of the Human Papillomavirus, Human Immunodeficiency Virus, and Oral Microbiota Interplay in Nigerian Youth (HOMINY) study^14^. Samples were processed in two sequencing batches: the first on the NovaSeq 6000 platform across three S4 flow cells (fc_2310, fc_2311, and fc_2401), and the second on the NovaSeq X platform on a single 25B flow cell (fc_b2; Methods). Functional profiles were inferred using two complementary strategies: (1) mi-faser^21, 22^, which annotated reads against the Fusion^20^ database, encompassing both characterized and unknown microbial functions, and (2) HUMAnN 3^23^, which generated EC number^25^ profiles, representing well-curated enzymatic functions. This large-scale, relatively deeply sequenced dataset provided a solid foundation to address two key questions: how much of the true functional diversity present in oral microbiome samples can be recovered by WMS, and how technical factors such as sequencing depth, flow cell, and annotation strategy shape that recovery.

### Microbial read depth, not total raw read depth, determines functional profiling power

A total of 671 youth oral rinse samples were sequenced (median=37.96 million raw read pairs per sample, [IQR: 31.16-43.99M]; **Table 1**) and quality filtered (median=35.46 million [IQR: 28.98-41.32M] per sample). Following host DNA removal, median of 7.27 million [IQR: 3.55-13.97M] microbial read pairs per sample remained, representing median=19% [IQR: 9-39%] of raw reads. While the numbers of raw and quality-filtered reads from each sample were, as expected, highly correlated (Pearson correlation coefficient (R)=0.97, Spearman correlation coefficient (ρ)=0.95), neither correlated well with the number of microbial reads detected (quality-filtered vs. microbial reads: Pearson R=0.14, Spearman ρ=0.02), reflecting substantial variability in host DNA contamination across samples.

**Table 1.** Sequencing yields and functional annotations across flow cells.

|  | fc_2310 | fc_2311 | fc_2401 | fc_b2 | All samples |
| --- | --- | --- | --- | --- | --- |
| *Number of samples* | 182 | 182 | 180 | 127 | 671 |
| *Raw read pairs (M)\** | 35.09 | 36.81 | 42.60 | 32.02 | 37.96 |
| *QC-ed read pairs (M)\** | 33.79 | 33.96 | 41.29 | 27.95 | 35.46 |
| *Microbial read pairs (M)\** | 10.48 | 9.92 | 5.09 | 5.88 | 7.27 |
| *Fusion functions detected\** | 24,158 | 24,257 | 21,826 | 21,806 | 22,928 |
| *Recovery of estimated Fusion function repertoire (%)\** | 62.42 | 62.22 | 57.26 | 55.63 | 60.26 |
| *EC numbers detected\** | 1,299 | 1,302 | 1,253 | 1,299 | 1,287 |

\*Medians for each flow cell and across all 671 samples.

This observation partially informed the next: the number of inferred Fusion functions (median=22,928 [IQR: 19,511-25,900]) was strongly correlated with microbial read counts (Pearson R=0.76, Spearman ρ=0.86), but not with the raw (Pearson R=0.14, Spearman ρ=0.05) or quality-filtered reads (Pearson R=0.16, Spearman ρ=0.04).

Because varied host DNA contamination decoupled raw read depth from microbial read depth, and microbial reads served as the input for all further functional annotation processes, all subsequent analyses were based on microbial read depth as the relevant measure of sequencing coverage for functional profiling.

### Microbial read depth is not linearly related to recovered functional diversity

In our 671 oral rinse samples, the median per sample microbial read pair count was 7.27 million [IQR: 3.55-13.97M]. Of these, 69% [IQR: 67-71%] mapped to the Fusion database, recovering a median of 22,928 [IQR: 19,511-25,900] functions per sample. Across all samples, the union of distinct Fusion functions comprised 69,702, with a median prevalence of each function across 49 [IQR: 3-474] samples (**SOM_Fig. 3A**).

To evaluate how sequencing depth shapes functional recovery, we mimicked shallower sequencing by randomly subsampling without replacement 10%, 30%, 50%, 70%, and 90% of microbial read pairs from each sample (Methods). Compared to the functions detected at each sample’s full microbial depth, the median fractions of functions retained were 51% [IQR: 44–56%], 74% [69–77%], 85% [82–87%], 92% [90–93%], and 98% [97–98%], respectively (**Table 2**). Thus, although functional diversity increased with sequencing depth, the rate of diversity gain diminished at higher depths, indicating a nonlinear relationship between sequencing depth and functional recovery.

**Table 2.** Retention of Fusion functions across lower microbial read depths.

| Subsampled microbial reads (% of full depth) | Median number of Fusion functions | Proportion of functions retained (%) of full-depth sample* | Proportion of functions retained (%) of total estimated repertoire** |
| --- | --- | --- | --- |
| 10 | 11,572 | 50.55 | 29.62 |
| 30 | 16,856 | 73.65 | 43.87 |
| 50 | 19,473 | 84.88 | 51.00 |
| 70 | 21,160 | 92.27 | 55.57 |
| 90 | 22,441 | 97.71 | 58.86 |
| 100 (full depth) | 22,928 | 100 | 60.26 |

\*median fraction of functions detected in subsampled read sets relative to the number recovered at full depth for each sample. \*\*median fraction of functions detected in subsampled read sets relative to the per-sample Hill curve estimate of the total functional repertoire.

Functions lost at reduced depth were predominantly those with low relative abundance in the original samples. For example, at 70% subsampling, 60% (median) of the functions from the lowest (10%) abundance bin of the original sample were lost; compared to 14% in the second lowest (10-20%) bin, and less than 1% in each of the higher-abundance (>20%) bins (**Figure 2A**).

**Figure 2:**
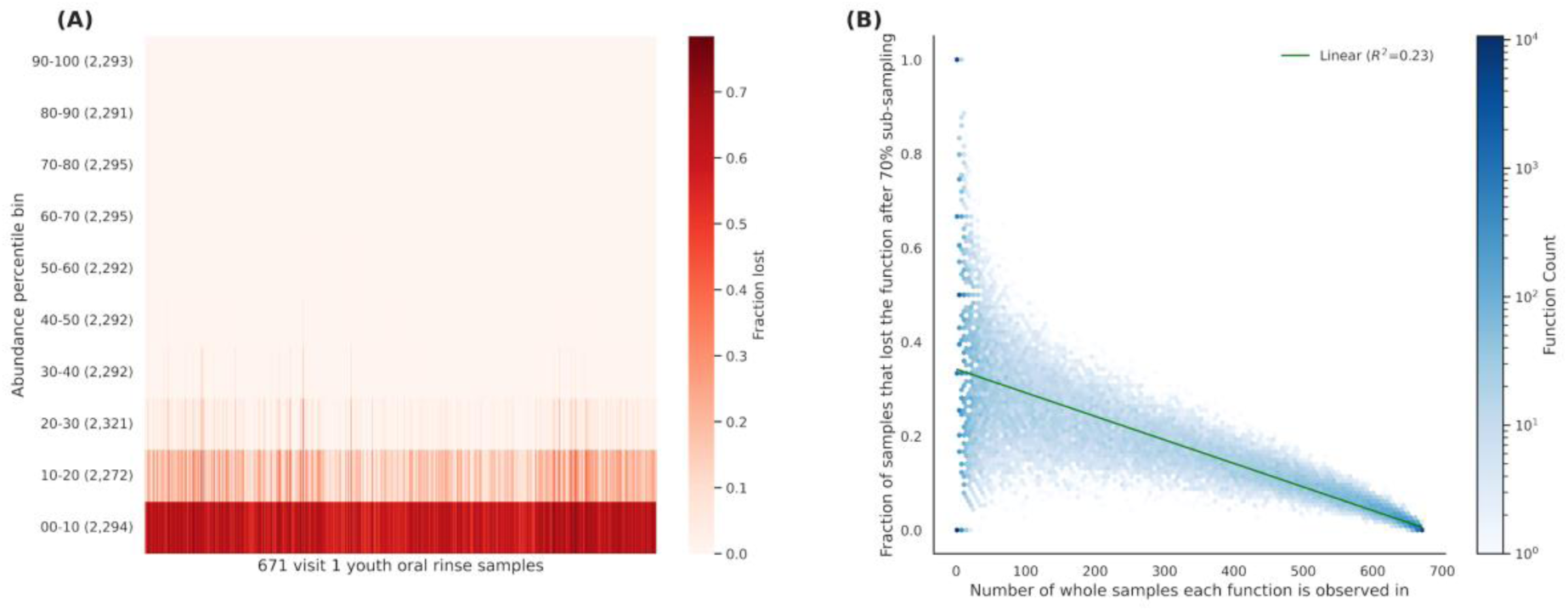
Low-abundance and low-prevalence Fusion functions are most vulnerable to dropout under reduced sequencing depth. **(A)** Heatmap showing the fraction of Fusion functions lost after down-sampling to 70% of the original microbial read depth across 671 oral rinse samples (x-axis). Functions are stratified by within-sample abundance deciles (y-axis), with the strongest dropout observed in the lowest abundance bins (0–10%), while higher-abundance functions are largely retained. **(B)** Relationship between Fusion function prevalence (x-axis; number of samples in which a function was observed in the full depth sequencing data) and dropout risk (y-axis; fraction of those samples in which the function was lost at 70% depth). Colour gradient indicates the density of functions reported within a particular location in the plot.

Function prevalence across samples was inversely correlated with dropout (Pearson R= 0.48, Spearman ρ= 0.25; **Figure 2B**). Thus, after 70% sub-sampling (30% microbial read reduction), a function present across ten samples at full depth could be absent from as many as eight (80%), but most often (median) from three (30%) of the samples. On the other hand, a function present in 300 samples at full depth, would most often only be lost from 17% of the samples (median=52, [IQR: 29-83]) after 70% sub-sampling.

Of 69,702 Fusion functions detected at full depth of sequencing across all samples, 7,376 (11%) were lost from more than half of their original samples at 70% sub-sampling. At full sequencing depth, these lost functions were most often found in only one sample (median=1, [IQR: 1-5]), compared to the median=49 samples [IQR: 3-474] prevalence for functions overall.

### Sequencing required for fully describing functional diversity is much deeper than expected

We modeled the relationship between microbial read depth and the number of detected Fusion functions to estimate the additional depth required for complete function recovery from a given sample. We compared three saturation curves: Michaelis-Menten (MM)^26, 27^, Weibull growth^28^, and Hill equation^29, 30^, to find the best fit across all simulated sequencing depths from the 671 samples. The Hill equation model provided the best fit in pooled curve comparisons (ΔAIC_(MM-Hill)_=981; ΔAIC_(Weibull-Hill)_=37) and also displayed the smallest residuals in per-sample fit, hold-out validation (leave-top-depth-out and leave-bottom-depth-out; Methods; **SOM_Table_1, SOM_Fig. 1-2**). We thus used the Hill model for all downstream analyses, including: (i) a pooled fit to all samples across all read depths (pooled Hill curve), (ii) per-sample fits at all depths within each sample, and (iii) sequencing depth planning (**Figure 3**).

**Figure 3:**
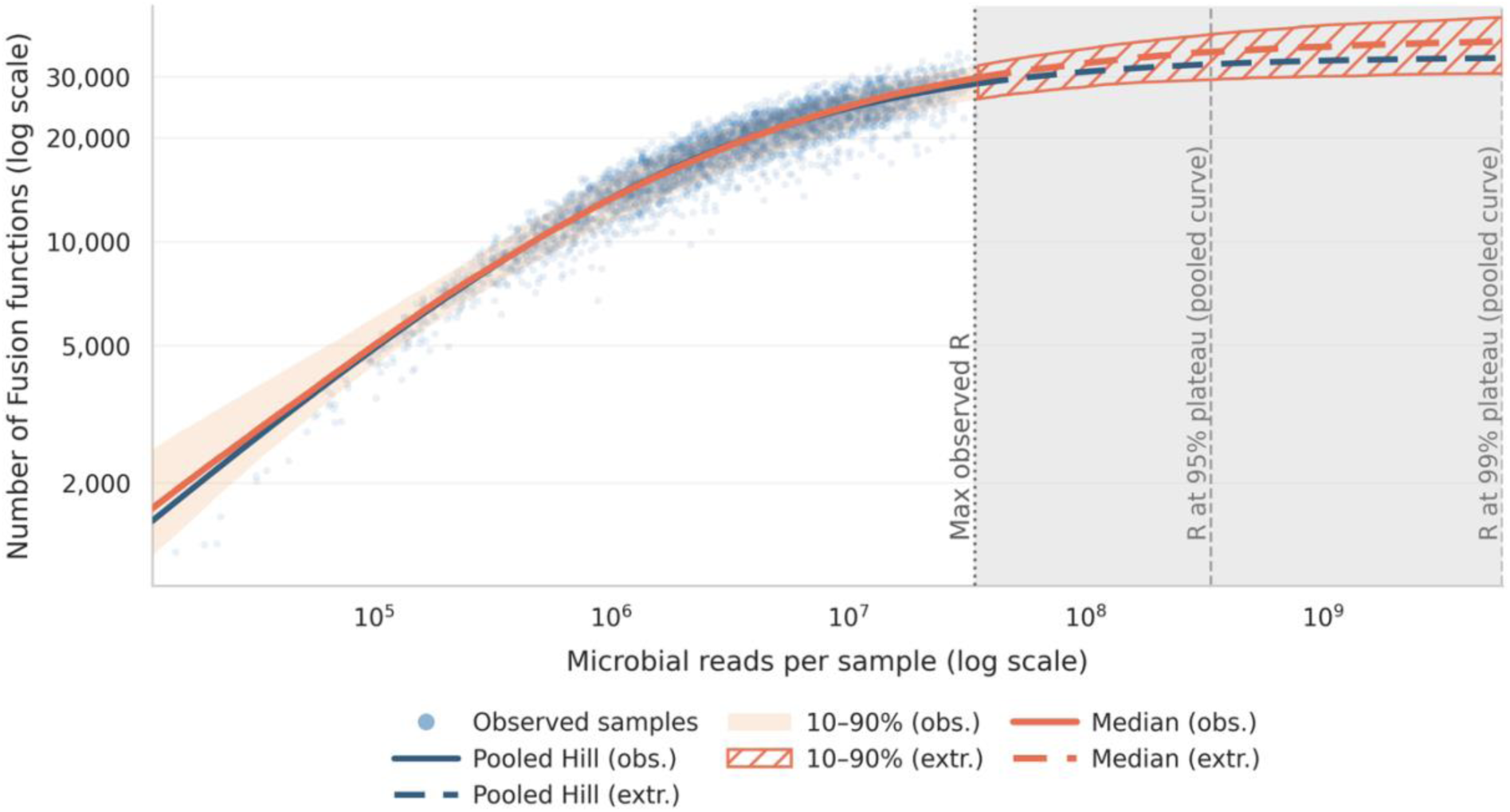
Fusion function detection across microbial read depths. Observed samples (blue dots) reflect the relationship between the microbial read depth and the number of detected Fusion functions. The pooled Hill model (solid blue line) was fit using all depths (10%, 30%, 50%, 70%, 90% sub-sampling and full depth) across 671 oral rinse samples. Per-sample Hill fits are summarized by the orange median curve with 10–90% variability bands. Extrapolation (dashed lines and shaded region) estimates the plateau of recoverable function and the corresponding microbial depths required. The leftmost vertical dashed line (Max observed R) marks the sample with the highest microbial read depth, while the two following dashed lines indicate the predicted microbial read depths required to achieve 95%, and 99%, respectively, of functions at plateau, as estimated by the pooled model.

Extrapolation of the pooled Hill curve (**Eqn. 3**; estimated parameters: plateau V_max_=34,331 functions, half-saturation constant K=2.18M microbial read pairs, and steepness h=0.58) indicated that 95% of the V_max_ plateau, i.e. 95% of all microbial functions truly present in the sample, could only be recovered at a depth providing 338M microbial read pairs (**SOM_Table_3**), a much higher sequencing depth than we strived for. At our actual full microbial read depths (median 7.3 M [IQR: 3.6-14.0 M] microbial read pairs per sample), per-sample function recovery was estimated to be only 60.26% [IQR: 51.88-66.30%]; estimated by per-sample Hill curves with parameters: V_max_=38,597 [IQR: 34,774-42,318], K=3.29M [IQR: 1.97-5.9M], and h=0.54 [IQR: 0.49-0.61]. These findings are consistent with a long tail of rare functions, unique to each sample.

Our benchmarks provide a quantitative basis for selecting sequencing depths to balance cost and function recovery and can be paired with function retention thresholds (**Table 3, SOM_table_3**) for study design.

**Table 3.**
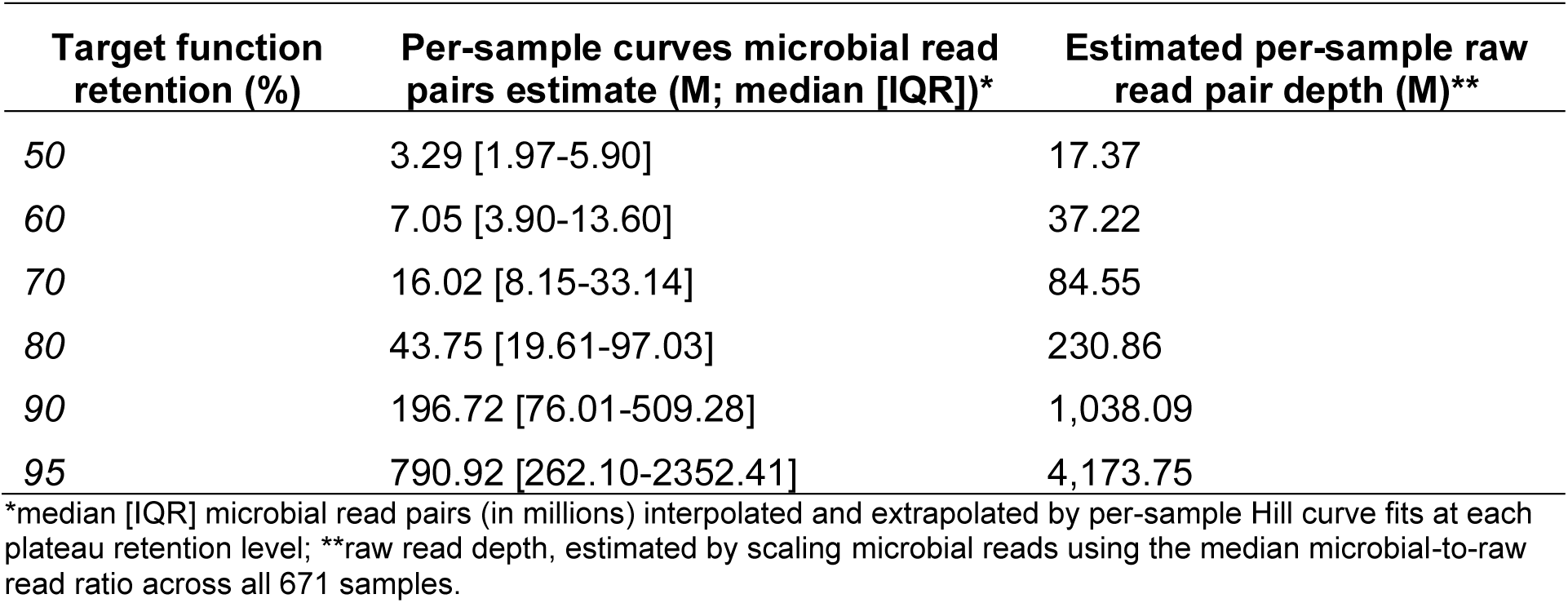
Estimated read depths necessary for achieving target fractions of total microbiome functionality.

| Target function retention (%) | Per-sample curves microbial read pairs estimate (M; median [IQR])* | Estimated per-sample raw read pair depth (M)** |
| --- | --- | --- |
| 50 | 3.29 [1.97-5.90] | 17.37 |
| 60 | 7.05 [3.90-13.60] | 37.22 |
| 70 | 16.02 [8.15-33.14] | 84.55 |
| 80 | 43.75 [19.61-97.03] | 230.86 |
| 90 | 196.72 [76.01-509.28] | 1,038.09 |
| 95 | 790.92 [262.10-2352.41] | 4,173.75 |

\*median [IQR] microbial read pairs (in millions) interpolated and extrapolated by per-sample Hill curve fits at each plateau retention level; \*\*raw read depth, estimated by scaling microbial reads using the median microbial-to-raw read ratio across all 671 samples.

### Function annotation strategy impacts sensitivity to down-sampling

To assess how function annotation strategy influences both (1) the fraction of microbial reads assigned to a function and (2) strategy sensitivity to sequencing depth, we compared functional profiles derived from full-depth and 70% subsampled microbial reads using two annotation strategies: Fusion database functions inferred by mi-faser and EC numbers inferred by HUMAnN 3 (Methods).

At full read depth, a median of 69% [IQR: 67-71%] of microbial reads were mapped to 22,928 [IQR: 19,511-25,900] Fusion functions per sample. However, only 13% [IQR: 12-14%] of the reads per sample were annotated with EC numbers (median 1,287 EC numbers per sample [IQR: 1,214-1,344]). When sequencing depth was reduced by 30% (70% sub-sampling), the number of Fusion functions declined by 7.73% [IQR: 6.65-9.52%] per sample. In contrast, EC annotations showed little change (1.00%), with 98.97% [IQR: 98.39-99.26%] of original EC functions retained.

Functions obtained using the two annotation strategies also showed distinct prevalence patterns (**SOM_Fig. 3**). At full depth, (a union of) 2,012 EC numbers were detected across all 671 samples, each present in a median of 621 samples [IQR: 48-669]. In contrast, 69,702 distinct Fusion functions were observed, but each was present in a median of only 49 samples [IQR: 3-474].

These observations highlight the coverage/recall tradeoff, i.e. broader coverage of rare, sample-specific functions by Fusion and the narrower, but consistent detection of EC numbers across more individuals. This further explains why Fusion-based annotations are more sensitive than EC numbers to reductions in read depth.

### Flow cell effect is significant for functional profiles

To test whether sequencing flow cells introduced systematic differences in profile functional compositions, we performed permutational multivariate analysis of variance (PERMANOVA) on Fusion-annotated functional profiles from samples distributed across the four flow cells. To better isolate the flow cell effects and minimize clinical variation, analyses were restricted to the 226 healthy subjects (NovaSeq 6000 S4 flow cells: fc_2310 n=22, fc_2311 n=90, fc_2401 n=59; NovaSeq X 25B flow cell: fc_b2 n=55; total N=226). The scale of profile differences was summarized using the pseudo-F statistic, which reflects the relative separation of groups compared with within-group variation. Overall, flow cell assignment explained a significant part of variation in functional profiles (abundances, Bray–Curtis dissimilarity pseudo-F=11.67, p=0.0001, R^2^_(flow cell)_=0.14; presence/absence, Jaccard dissimilarity pseudo-F=6.11, p=0.0001, R^2^_(flow cell)_=0.08). That is, flow cell batch effects contributed significantly to the detectable differences between samples.

Pairwise PERMANOVA tests further highlighted significant differences in functional composition across all flow cells (Benjamini-Hochberg adjusted for each flow-cell pair; **SOM_Table_4**). Abundance-based (Bray-Curtis) comparisons between flow cells were all significant, with the largest difference observed between flow cell types *fc_2401* (NovaSeq 6000, cell type S4) and *fc_b2* (NovaSeq X, cell type 25B; pseudo-F=25.87, p_adj=0.00015, R^2^_(flow cell)_=0.19). Presence/absence (Jaccard) analyses yielded a similar outcome, with all pairwise comparisons significant and the greatest separation observed between flow cell types *fc_2311* (NovaSeq 6000 S4 cell) and *fc_b2* (NovaSeq X 25B cell; pseudo-F=9.95, p_adj=0.0002, R^2^_(flow cell)_=0.07).

Together, these results demonstrate that flow cell assignment significantly influenced functional annotation of samples. The strongest effects were observed between different sequencing platforms (NovaSeq 6000 S4 vs. NovaSeq X), but notable heterogeneity was also detected among flow cells within the same NovaSeq 6000 S4 platform. This highlights the importance of accounting for flow cell assignment in downstream analyses to avoid confounding technical effects with biological variation.

## Discussion

Here, we developed a framework for relating the whole metagenome shotgun sequencing (WMS) read depth with recovery of microbial functions. Using our models we were able to estimate the full functional potential of the oral microbiomes in our data. We then systematically evaluated how technical factors, including sequencing depth, annotation strategy, and flow cell batch effects can influence the recovery of microbial functions from WMS data. Below we summarize our findings for the analysis of our cohort and suggest that these could elucidate steps/directions for similar types of research.

### Experimental design is disconnected from attainable microbial function annotations

Our analysis of WMS data was complicated in the experimental design phase by two critical disconnects between the planned sequencing depth of an experiment and the microbial functional information ultimately obtained.

First, targeted vs. realized raw sequencing depth are drastically different. In our first sequencing batch (NovaSeq 6000 S4), we targeted 44.4-55.6 million paired-end reads per sample based on the flow cell capacity and the number of samples on each cell. The realized raw read yields after demultiplexing were lower and variable across the three flow cells, with medians of 35.1M, 36.8M, and 42.6M reads per sample, as much as a fifth of expected reads missing.

Such deviations from ideal are common in multiplexed runs, where per-sample output depends on run quality, variation in per-sample DNA input during pooling, and routine losses. Sequencing facilities therefore commonly budget ±20% variability in delivered reads relative to the target depth^31, 32^. Our realized median per-sample yields corresponded to 79%, 83%, and 96% of the minimum read target across the three S4 flow cells. This gap highlights that “targeted depth” at the design stage does not guarantee the same realized depth in sequencing output. In studies where a strict minimum actual depth per sample is essential, e.g. metagenome-assembled genomes recovery^33-36^, it may be necessary to budget an additional safety margin above the target depth, considering all factors influencing the per-sample expected read counts.

Second, raw read depth is not indicative of microbial read depth. Sequencing depth is a foundational consideration for any WMS study design; it is typically measured and reported as the total number of raw read pairs generated per sample^7, 37^. However, in many host-associated microbiomes, this convention is misleading as host DNA contamination varies widely across samples. Reported host DNA content ranges from 61-99% in vaginal samples^38, 39^, 57-94% in nasal samples^40, 41^, and 60-90% in subgingival plaque^42^. In our oral rinse samples, microbial reads accounted for only 19% [IQR: 9-39%] of the raw reads, with an additional median 4% [IQR: 3-12%] of raw reads discarded during quality control steps.

It is the microbial read depth, i.e. the number of quality-filtered reads remaining after host DNA removal, however, rather than raw read counts, that is the primary determinant of functional richness in WMS profiles. Consequently, planning sequencing efforts based solely on raw read counts is difficult, particularly problematic in low-microbial-biomass environments^38, 43, 44^ (e.g. oral, vaginal, skin), as variable host DNA contamination decouples raw sequencing output from usable microbial signals.

Variation in host DNA content can arise from technical factors, such as sample collection, storage, and DNA extraction methods^11, 12, 38, 45-47^. Biological factors not being considered in the specific study, such as site-specific physiology or infection, can also contribute^45, 48^. Host DNA depletion strategies prior to sequencing, such as enzymatic digestion, methylation-based removal, physical separation, or chemical depletion, can be considered to enrich microbial content^8, 11, 38, 45^. Overall, unless host DNA variation itself is biologically informative, it represents technical noise that can potentially confound analyses of microbial functions.

### Recovering the full microbial functional repertoire requires much more sequencing depth than typically achievable

To assess the breadth of functionality captured at different depths, we performed per-sample subsampling of microbial reads from the original dataset at five levels (10%-90%, 20% increments; **Table 2**). We then quantified the fraction of functions retained relative to those detected at each sample’s full depth. Although function discovery rates slowed down with increasing depth, e.g. increasing from 10% to 30% of reads yielded 5203 (median) more functions, whereas increasing from 70% to 90% yielded only 1256 more, novel functions continued to emerge at all depths (**Table 2**). This indicates that even our full-depth data underestimates the true functional repertoire.

Across the 671 original oral rinse samples, we detected 69,702 unique Fusion functions. Of these, only 2,308 (3%) were present in every sample, while the majority followed a long-tail distribution (median prevalence: 49 samples; IQR: 3-474; **SOM Fig. 3A**). Curiously, this number is much higher than the 344 EC numbers common to all samples. Subsampling further revealed that functions most likely to disappear at reduced sequencing depths were primarily derived from this long tail; i.e. those with low prevalence across the cohort. We also note that functions with low relative abundance within samples were often lost (**Figure 2A-B**). Thus, even at current full sequencing depths, uncaptured functions likely represent the rarer and less abundant components of the functional landscape. We suspect that these, however, are often prime candidates for establishing host phenotypic differences.

Previous studies have proposed modeling frameworks to extrapolate the relationship between sequencing depth and information recovery^49-52^. For example, Hooper *et al* (2010) modelled genomic coverage as Poisson-distributed read counts per bin, with bin expectations drawn from a Gamma distribution to account for bin abundance differences^49^. Monleon-Getino & Frias-Lopez (2020) adapted ecological rarefaction calculation approaches to simulated meta-transcriptome profiles, fitting Weibull growth models to estimate asymptotic gene recovery^50^. These approaches, however, were mostly developed using mock microbial communities with simulated reads or microbial profiles under specific assumptions. Thus, they did not directly align with our goal of characterizing the complete functional potentials of real oral microbiomes. We therefore evaluated multiple candidate saturation models widely used in ecology^50, 53, 54^, ultimately selecting the Hill equation for its superior fit to our data.

Hill curve extrapolations suggested over 38,597 Fusion functions exist per sample, and that 3.29M microbial reads are required to recover half of this repertoire. Thus, at our actual median depth of 7.3M microbial reads per sample, only 60% of the predicted functional potential was captured (**Table 2**). By contrast, even 10% of our reads were sufficient to recover 30% of all putative functions, reflecting the strong representation of common and abundant functions at shallow depths. Achieving 90% functional recovery, however, would require 197M microbial reads per sample (converts to 1.04 billion raw read pairs; **Table 3**), a depth that is impractical for large-scale studies.

Note that we did not apply bootstrap resampling to further quantify down-sampling uncertainty. The reported medians and percentile ranges across fitted curves may therefore further underestimate true variability of sample depth to function mappings.

Our findings highlight the inherent difficulty of capturing the complete microbiome functional repertoire using sequencing at field-standard depths. This limitation is further compounded by the fact that individual or group-specific functions are disproportionately at risk of dropout, a challenge we explore below.

### Low-prevalence and group-specific functions are more likely to disappear at lower sequencing depth

Function prevalence across samples, i.e. the number of samples in which a function is detected, was a strong predictor of dropout risk in down-sampling. Functions with low prevalence were particularly prone to loss. For example, after a 30% reduction in microbial reads, functions originally detected in ten individuals were likely absent in a third of those samples, whereas functions detected in 300 samples were dropped from fewer than a fifth of profiles (**Figure 2B**).

This impact of sequencing depth on prevalence of detectable functions has important implications for the design and interpretation of microbiome studies. In clinical or environmental-shift microbiome studies, sample groups are often imbalanced. Functions specific to smaller groups, e.g. disease cases, are, by definition, represented in fewer samples and, unless very abundant within those samples, face a higher risk of dropout with shallow sequencing. This challenge can be amplified in pooled or multiplexed sequencing designs, where functions confined to a small subset of samples contribute little to the total library and may still be lost even if not rare within their subgroup. This creates a potential source of systematic bias, where group-specific functional signatures may be underrepresented and, thus, unexplored.

We suggest that experimental design should be considered alongside sequencing depth planning, with attention to both adequate sample representation and sufficient microbial read depth. Together, these factors will help ensure more robust detection of functional signals, even for functions restricted to smaller groups within the cohort.

### Trade-offs between function coverage and recall across annotation strategies

To evaluate how annotation strategy influences functional recovery and depth-independent stability, we compared Fusion functions with EC numbers. Earlier studies show that functional databases differ in coverage, gene grouping, and granularity, substantially influencing both the proportion of reads that can be classified, and the resulting function profiles and their downstream interpretation^6, 7, 55^. Similarly, we observed that while Fusion annotated a median of 69% of microbial reads (a median of 22,928 Fusion functions per sample), EC only captured 13% of the reads(a median of 1,287 ECs per sample). This contrast reflects the distinct philosophies of the two approaches: Fusion maximizes breadth by capturing both known and uncharacterized functions, while EC numbers emphasize well-curated enzymatic activities.

Indeed, at full sequencing depth, Fusion functions typically exhibited lower prevalence across individuals (median 49 samples; **SOM_Fig. 3A**) than most EC numbers, which were highly prevalent (621 samples; **SOM_Fig. 3B**). The high prevalence of nearly all enzymatic functions in our cohort, i.e. 344 out of the total 2,012 ECs present in every sample, suggests that, within the context of EC annotations, phenotypic differences are likely to be primarily captured by functional abundance than by presence/absence metrics. Note that 2,308 Fusion functions were identified across all samples in our cohort, a number higher than what was captured by EC annotations. While we did not explicitly evaluate the amount of overlap between these functions, in the very least this discrepancy suggests that Fusion was able to annotate seven times more common bacterial functionality than that accounted for by enzymes alone.

When sequencing depth was reduced by 30% (i.e. 70% sub-sampling of microbial reads), the number of Fusion functions dropped by 8% vs the original number of functions in that sample, while the number of ECs was minimally affected (1% lost). These results indicate that databases with broader functional coverage and higher read-mapping rates, such as Fusion, are more sensitive to reductions in sequencing depth because they include many functions supported by few reads or restricted to few samples. In contrast, curated sets involving specific well-studied function groups like EC numbers capture fewer but more consistently observed functions, making them more robust to variation in sequencing depth at the cost of excluding rare and uncharacterized signals.

Together, these findings highlight a trade-off between annotation breadth (function coverage) and stability (function recall). Decisions about sequencing depth and annotation strategy should therefore be considered in light of study goals: if the focus is on well-studied, common molecular functions, moderate sequencing depths may be sufficient for deep coverage. Conversely, if the goal is to explore the full functional repertoire including rare or uncharacterized functionality, substantially deeper sequencing will be required. In our cohort, even at full sequencing depth (7.3M microbial read pairs; 38M raw read pairs per sample), only 60% of the projected Fusion functional repertoire per sample was estimated to have been recovered, underscoring the challenge of capturing the functional landscape in its entirety.

We note that while our work reported here has focused on the oral microbiome, we believe that our models can be similarly applied to explore function extraction from sequencing of a wide range of host associated, and possibly environmental, microbiomes.

## Conclusions

Here we explored the microbiome extracted from the oral rinse samples of Nigerian youths. We developed a framework to link read depth to functional recovery, providing a practical tool to guide sequencing decisions and balance cost against the scope of functions targeted. Using our models, we found that a large fraction (40%) of microbial function diversity in these microbiomes remained undetected even at sequencing depths of over 38 million read pairs per sample. We further report that fully recovering the complete microbial function repertoire from metagenomic data may require much more sequencing than expected.

We note that the choice of annotation strategies in this work also influenced how sequencing depth translated into functional information: Fusion covered a wider range of functions but was highly sensitive to depth shifts, whereas EC numbers yielded more stable profiles at the cost of excluding rare and uncharacterized signals. Finally, flow cell differences introduced systematic shifts in functional composition, underscoring the importance of accounting for batch effects in large-scale studies.

Together, these findings advance our understanding of the limits of functional discovery in oral microbiomes, and offer concrete principles for future study design. Integrating sequencing depth planning with annotation choice and careful batch control will be essential to minimize technical bias and improve reproducibility and interpretability of microbiome research across host-associated environments.

## Declarations

### Ethics approval and consent to participate & Consent for publication

This work used already generated data from participants as described in (https://doi.org/10.1136/bmjopen-2024-091017), from which ethics approval and consent can be found.

### Availability of data and material

Microbial reads after low quality and host DNA removal: BioProject PRJNA1345363. Fusion and EC functional profiles: https://figshare.com/s/c4e7290ede75464f70a1.

### Competing Interests

Dr. Osazuwa-Peters was a scientific advisor to *Navigating Cancer* and has received consulting fees from *Merck* for consultation on HPV vaccination.

### Funding

The work of Y.B., J.L., MO.C., N.O-P., O.P., NL.I., N.F.S., O.O., FE.E. was supported by grant R01-DE032216 from the National Institute of Dental and Craniofacial Research.

### Authors’ contributions

O.P., NL.I., O.O., FE.E.: collected and processed samples used in this study. J.L., Y.B.: implemented data analysis, model fitting, and wrote the main manuscript. MO.C., N.F.S., N.O-P., O.P., and remaining authors reviewed and provided helpful suggestions to the manuscript.

## Acknowledgements.

We are grateful to R. Prabakaran and Jay Mayfield for all helpful discussions and to Sergio Gramacho and Aiden Maloney-Bertelli for technical support (all Emory University). We truly appreciate all study participants and their families for their participation in the study that is the basis of this work. We thank all members of the HOMINY study team at the University of Benin Teaching Hospital, Nigeria, as well as the nurses, doctors, and research staff of the Nigerian labs who have facilitated data collection for this study. We thank Aaron Ericsson, Giedre Turner, Rebecca Dorfmeyer, and Nathan J. Bivens at the University of Missouri Genomics Technology Core for their sequencing service. We also thank all those who deposit their experimental results into public databases and those who maintain these databases.

## Notes

https://figshare.com/s/c4e7290ede75464f70a1

