## Supplemental Material for "Whole Metagenome Sequencing: not Deep Enough for Complete Microbial Function Recovery"

**This file includes:**

Figs. S1 to S3
Tables S1 to S4

### Supplementary Figures

#### Fig. S1

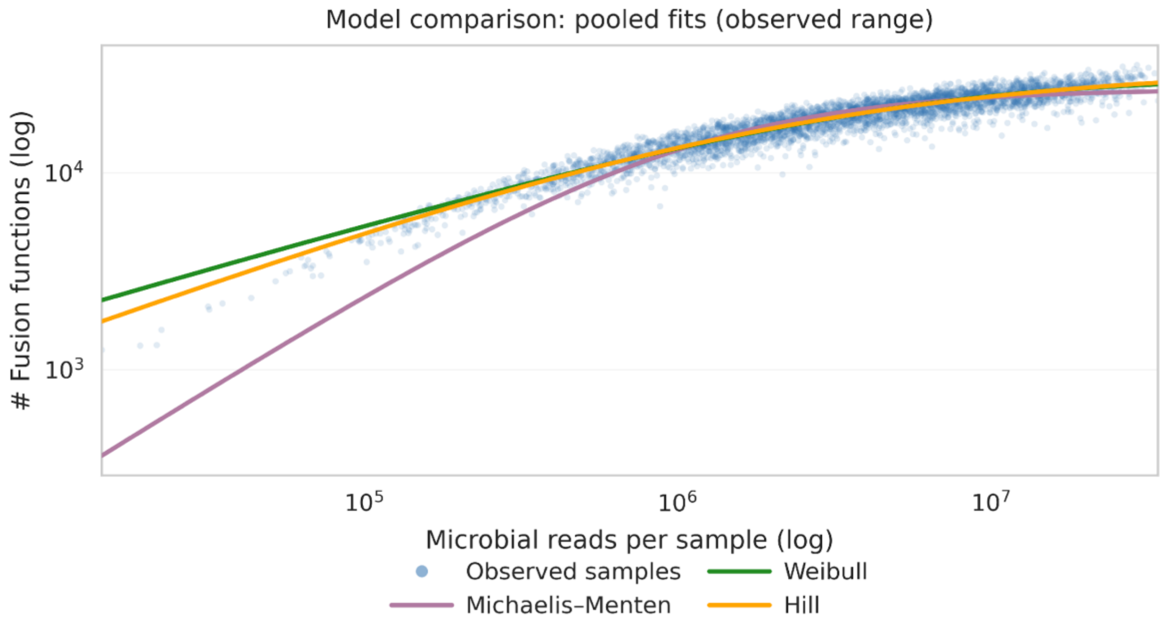


**Fig. S1 Candidate saturation models fit to pooled microbial depths - Fusion function counts data.** Pairs of microbial read counts and corresponding inferred Fusion function counts were pooled across all 671 oral rinse samples and microbial read depths (10%, 30%, 50%, 70%, 90% subsampled, and full depth). Three candidate models: Michaelis-Menten, Weibull, and Hill equation were fit to the pooled dataset to evaluate their ability to capture the observed relationship between microbial depth and Fusion function recovery.

### Fig. S2

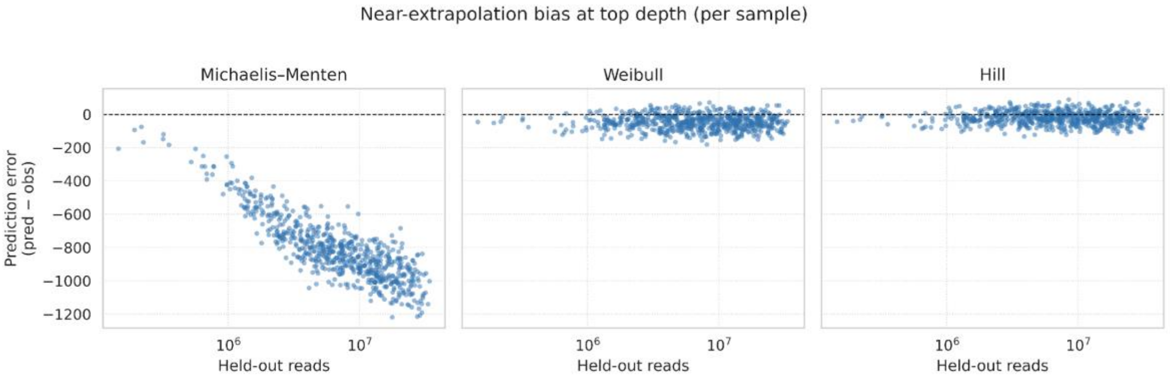


Fig. S2 (A)

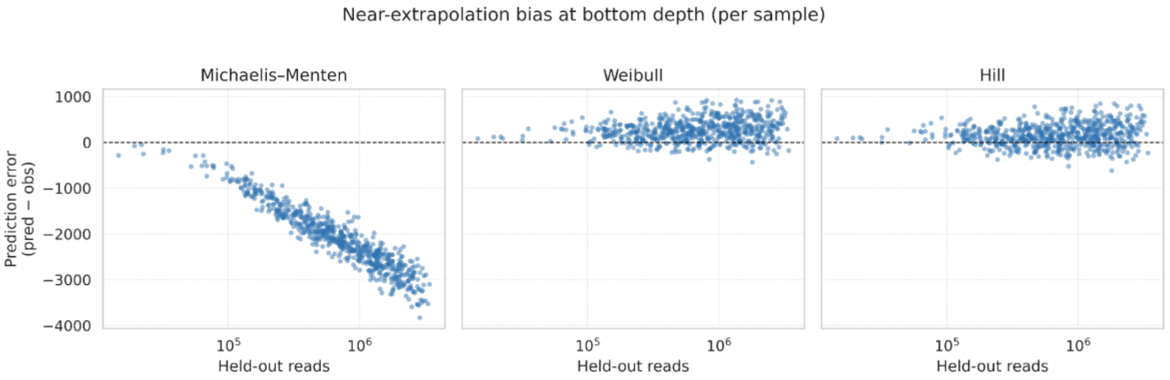


Fig. S2 (B)

**Fig. S2 Prediction error of candidate saturation per-sample models on held-out depths.**
Prediction accuracy of the Michaelis-Menten, Weibull, and Hill models was evaluated by per-sample leave-one-depth-out validation.
(A) *Top-depth test:* For each model, per-sample curves were fit using 10%, 30%, 50%, 70%, and 90% sub-sampled microbial read pairs and their corresponding Fusion function counts. The fitted curves were then used to predict function counts at the left-out full microbial read depth. Plots show the relationship between per-sample full-depth microbial read pairs and prediction error (predicted minus observed function count).
(B) *Bottom-depth test:* For each model, per-sample curves were fit using 30%, 50%, 70%, 90%, and full microbial read depths. The fitted curves were then used to predict function counts at the left-out 10% depth. Plots show prediction error as in panel A.

### Fig. S3

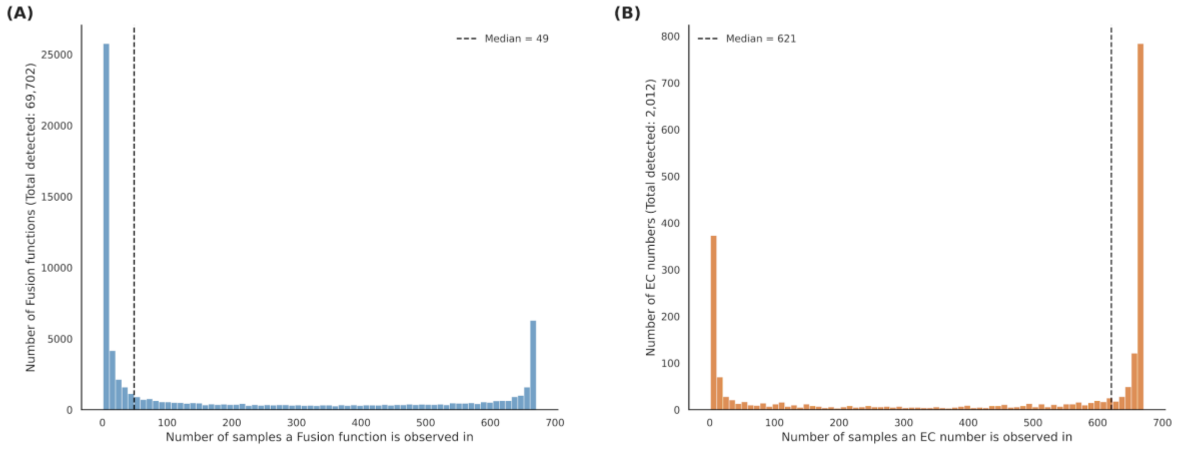


**Fig. S3 Prevalence distribution of Fusion functions and EC numbers across 671 oral rinse metagenomes.**
(A) Histogram showing the number of samples in which each Fusion function was detected (total detected: 69,702). The median prevalence was 49 samples. (B) Histogram showing the number of samples in which each EC number was detected (total detected: 2,012). The median prevalence was 621 samples. Vertical dashed lines indicate median values.
.

**Table S1. Model performance metrics for fitting Fusion function saturation with microbial read depth.**

| Model | Number of Params | $R^2$ | RMSE | MAE | AIC | BIC |
|---|---|---|---|---|---|---|
| ***Michaelis–Menten*** | 2 | 0.858 | 2257.278 | 1787.616 | 62180.860 | 62193.461 |
| ***Weibull growth*** | 3 | 0.887 | 2007.095 | 1539.826 | 61236.981 | 61255.882 |
| ***Hill*** | 3 | 0.888 | 1997.899 | 1528.776 | 61200.004 | 61218.906 |

Models were fit to pooled data relating microbial read depth to Fusion functions detected across 671 oral rinse samples (10%, 30%, 50%, 70%, 90% subsampled and full depth). Metrics reported include coefficient of determination ($R^2$), root mean squared error (RMSE), mean absolute error (MAE), and information criteria (AIC, BIC).

**Table S2. Sequencing yields and functional annotations across four flow cells (oral rinse samples from 226 healthy participants).**

| | fc_2310 | fc_2311 | fc_2401 | fc_b2 |
|---|---|---|---|---|
| *Number of healthy samples* | 22 | 90 | 59 | 55 |
| *Raw read pairs (M)* | 35.30 | 37.91 | 43.14 | 37.22 |
| *QC-ed read pairs (M)* | 34.29 | 34.00 | 41.64 | 34.27 |
| *Microbial read pairs (M)* | 15.24 | 10.06 | 5.27 | 5.88 |
| *Number of Fusion functions* | 24,852 | 24,675 | 21,890 | 21,590 |

Medians for healthy samples within each flow cell. Metrics include sequencing depth (raw, QC-passed, and microbial read pairs) and functional annotations (Fusion functions).

#### Table S3. Estimated microbial read depth required to achieve target fractions of plateau functions.

| Target plateau retention (%) | Pooled model estimate (M read pairs) | Per-sample curves 10% quantile estimate (M) | Per-sample curves median estimate (M) | Per-sample curves 90% quantile estimate (M) |
|---|---|---|---|---|
| *50* | 2.18 | 1.24 | 3.29 | 10.74 |
| *60* | 4.36 | 2.28 | 7.05 | 25.83 |
| *70* | 9.29 | 4.38 | 16.02 | 67.62 |
| *80* | 23.41 | 9.75 | 43.75 | 243.26 |
| *90* | 93.97 | 33.39 | 196.72 | 1499.71 |
| *95* | 338.2 | 100.38 | 790.92 | 8090.66 |

Required microbial read pairs (in millions) to retain targeted fraction of functions were estimated from the pooled Hill model, with 10%, 50% (median), and 90% quantiles summarized from per-sample Hill curves estimations.

#### Table S4. Pairwise PERMANOVA results testing differences in healthy participants' functional composition across combinations of flow cells.

| Comparison pairs (flow cells) | N | Pseudo-F | R2 | Raw p-value | Adjusted p-value | Significant (FDR < 0.05) | Metric |
|---|---|---|---|---|---|---|---|
| **fc_2310 vs. fc_2311** | 112 | 2.58 | 0.0229 | 0.0148 | 0.0148 | Yes | Bray–Curtis |
| **fc_2310 vs. fc_2401** | 81 | 6.07 | 0.0713 | 0.0001 | 0.00015 | Yes | Bray–Curtis |
| **fc_2310 vs. fc_b2** | 77 | 3.58 | 0.0455 | 0.0014 | 0.00168 | Yes | Bray–Curtis |
| **fc_2311 vs. fc_2401** | 149 | 17.02 | 0.1037 | 0.0001 | 0.00015 | Yes | Bray–Curtis |
| **fc_2311 vs. fc_b2** | 145 | 10.39 | 0.0677 | 0.0001 | 0.00015 | Yes | Bray–Curtis |
| **fc_2401 vs. fc_b2** | 114 | 25.87 | 0.1877 | 0.0001 | 0.00015 | Yes | Bray–Curtis |
| **fc_2310 vs. fc_2311** | 112 | 1.71 | 0.0153 | 0.0326 | 0.0326 | Yes | Jaccard |
| **fc_2310 vs. fc_2401** | 81 | 3.87 | 0.0467 | 0.0005 | 0.00075 | Yes | Jaccard |
| **fc_2310 vs. fc_b2** | 77 | 4.16 | 0.0525 | 0.0010 | 0.0012 | Yes | Jaccard |
| **fc_2311 vs. fc_2401** | 149 | 8.31 | 0.0535 | 0.0001 | 0.0002 | Yes | Jaccard |
| **fc_2311 vs. fc_b2** | 145 | 9.95 | 0.0651 | 0.0001 | 0.0002 | Yes | Jaccard |
| **fc_2401 vs. fc_b2** | 114 | 5.41 | 0.0461 | 0.0001 | 0.0002 | Yes | Jaccard |

Pairwise PERMANOVA were performed based on Bray-Curtis (abundance-weighted) and Jaccard (presence/absence) dissimilarities. Reported values include the pseudo-F statistic, unadjusted and adjusted p-values (Benjamini–Hochberg correction), total number of samples compared (*N*), and significance status. Flow cell sample sizes were: *fc_2310* (n = 22), *fc_2311* (n = 90), *fc_2401* (n = 59), and *fc_b2* (n = 55).
